# Loss of cytoskeletal proteostasis links dysregulation of cell size and mechanotransduction in mesenchymal stem cell senescence

**DOI:** 10.1101/2022.10.09.511462

**Authors:** Venkatesh Mallikarjun, Oana Dobre, Mark R. Jackson, Melissa Kidd, Jack Llewellyn, Hamish T. J. Gilbert, Stephen M. Richardson, Joe Swift

**Affiliations:** Wellcome Centre for Cell-Matrix Research, University of Manchester, Manchester, M13 9PT, UK; Division of Cell Matrix Biology and Regenerative Medicine, School of Biological Sciences, Faculty of Biology, Medicine and Health, Manchester Academic Health Science Centre, University of Manchester, Manchester, M13 9PT, UK

**Keywords:** Mechanotransduction, Mesenchymal stem cells (MSCs), Senescence, Ageing, Proteostasis

## Abstract

Tissues are maintained by homeostatic feedback mechanisms where cells respond to, but also modify, the chemical and mechanical properties of the surrounding extracellular matrix. Mesenchymal stem cells (MSCs) resident in the marrow niche experience a diverse mechanical environment, but ageing can affect the composition and quality of bone and marrow tissues. Here we quantified the effect of replication-induced senescence on MSC morphology and their ability to correctly respond to different substrate stiffnesses. The matrix proteome was found to be sensitive to substrate stiffness, but pharmacological inhibition of cellular contractility perturbed this response, decreasing levels of tenascin-C, fibulins and fibronectin. Similar decreases in these mechanosensitive proteins were observed in senescent cells, suggested a decoupling of mechanotransduction pathways. Intracellular proteomic and transcriptomic analyses confirmed a decrease in components of the cytoskeletal chaperone complex CCT/TRiC in senescent MSCs. Finally, pharmacological inhibition of CCT/TRiC was able to partially recapitulate senescence-associated morphological changes in non-senescent MSCs. These results demonstrate a senescence-mediated perturbation to cytoskeletal homeostasis, pathways of mechanotransduction and the secretion of matrix proteins required for tissue maintenance.

## INTRODUCTION

Mesenchymal stem cells (MSCs) resident in bone marrow experience a diverse mechanical environment (Ivanovska *et al*, 2017) (Fig. 1A), but ageing can alter the composition and quality of bone and marrow tissues (Justesen *et al*, 2001), as well as cause changes to the physical loads experienced by ageing bones (Gurkan & Akkus, 2008). Mechanical stimulation has been shown to influence MSC behaviour, including control over transcript and protein levels, morphology, motility and lineage potential (Engler *et al*, 2006; McBeath *et al*, 2004; Raab *et al*, 2012; Swift *et al*, 2013). MSCs cultured on polyacrylamide (PA) hydrogels with stiffness commensurate with soft marrow were found to exhibit balled cellular and nuclear morphologies and favour adipogenesis; MSCs on stiffer hydrogels showed greater spreading, with correspondingly increased nuclear areas, and were more prone to osteogenesis (Swift *et al*., 2013). Furthermore, morphological responses to substrate stiffness established within hours and days have been shown to be predictive of lineage specification several weeks later (Treiser *et al*, 2010). Mechanically-induced morphological changes are known to propagate into transcriptional regulation through mechanotransduction pathways, such as the translocation of transcription factors (Connelly *et al*, 2010; Dupont *et al*, 2011), or transmission of mechanical stimulation through the cytoskeleton and nucleoskeleton to affect chromatin organisation (Tajik *et al*, 2016). Age-related changes in mechanical properties have been characterised in a range of tissues, including those such as skin, heart and lung where mechanics determine or limit functionality (Phillip *et al*, 2015; Sicard *et al*, 2018). However, it is not well understood how ageing affects the relationship between tissue mechanics and cell regulation.

**Figure 1.**
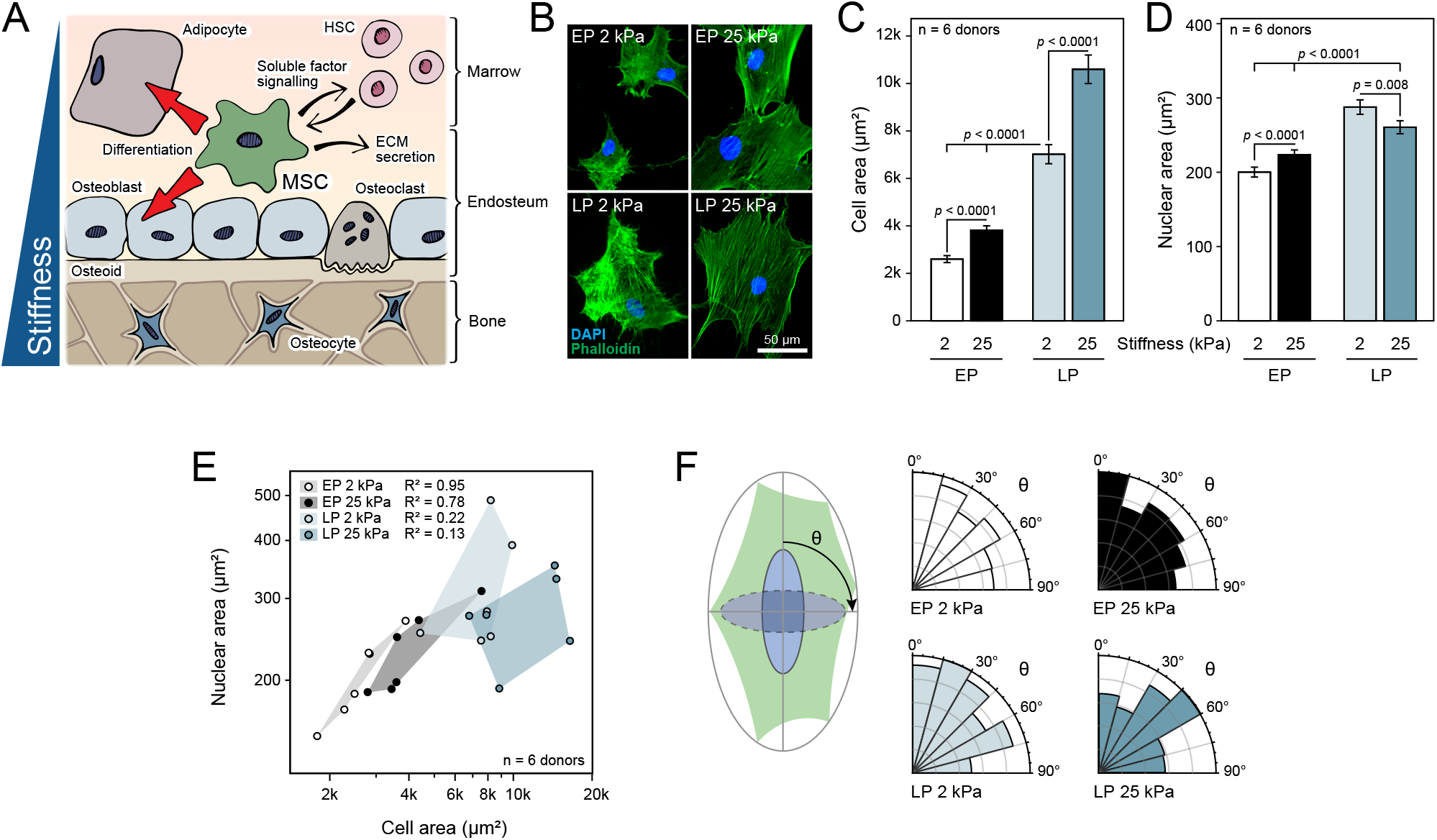
Senescence perturbs the coordinated response to substrate stiffness in cells and their nuclei. (**A**) Cartoon illustrating the roles of mesenchymal stem cells (MSCs) within the bone marrow niche. MSCs are responsive to the mechanically diverse properties of the niche and have been shown to favour adipogenesis and osteogenesis in soft and stiff microenvironments respectively (Ivanovska *et al*., 2017). Factors secreted by MSCs contribute to the extracellular matrix (ECM) and the maintenance of haematopoietic stem cell (HSC) populations (Anthony & Link, 2014). (**B**) Representative images (DAPI, blue; phalloidin, green) of early passage (EP, P3) and late passage (LP, P8) MSCs cultured on soft (2 kPa) or stiff (25 kPa) collagen-I coated polyacrylamide (PA) hydrogels for three days. (**C**) Quantitative analysis of cell spread area and (**D**) nuclear area of EP (P3 – 6) and LP (P6 – 12) MSCs cultured for four days on soft or stiff PA hydrogels. EP and LP MSCs spread to a significantly greater extent on stiffer substrates (*p* < 0.0001), although LP cells were consistently significantly larger (*p* < 0.0001). Nuclear area was correspondingly greater in EP MSCs on the stiff substrate (*p* < 0.0001), but this response was reversed in LP MSCs (*p* = 0.008). In figure parts (C) and (D), bar charts show mean ± SEM; significance was determined by F-tests on linear models including treatment and donor terms, *n* = 6 donors; individual donor distributions can be found in Supplementary Figs. S1H, I. (**E**) The relationship between mean cell and nuclear spread area was correlated in EP MSCs from multiple individual donors on both soft and stiff substrates (Pearson’s R-squared of 0.98 and 0.78, respectively; *n* = 6 donors). This correlated behaviour was diminished when the same cells were cultured to late passage. (**F**) Analysis of the orientation of the nucleus with respect to the longest axis of the cell in EP and LP MSCs on soft (2 kPa) and stiff (25 kPa) substrates. Alignment was significantly decreased in LP vs. EP MSCs on stiff substrates (*p* = 0.02 from ANOVA testing; *n* = 6 donors).

Stem cell dysfunction is regarded as a hallmark of ageing (Lopez-Otin *et al*, 2013) and plays a key role in the failure of tissues in elderly individuals to regenerate correctly following injury. Increased cellular senescence in stem cell pools is considered to be a major cause of stem cell dysfunction and subsequent regenerative failure in aged tissues. Cellular senescence is thought to have evolved as a defence mechanism against cancer (Collado & Serrano, 2010) and typically involves irreversible cell cycle arrest, thus preventing dysfunctional cells from proliferating. Senescent cells also serve an important function in mediating an inflammatory response to direct wound healing following injury (Chiche *et al*, 2017; Ritschka *et al*, 2017). However, in elderly individuals, a dysfunctional immune system allows senescent cells to linger, promoting the chronic “sterile” inflammation which has been described as a contributing factor in many age-related diseases (Acosta *et al*, 2013). Clearance of senescent cells that accumulate with age in mice has been shown to ameliorate the progression of age-related disease (Baker *et al*, 2016), demonstrating their causative role in these diseases.

Here we deployed multi-omic analysis to quantify the effects of substrate stiffness and replication-induced senescence on the primary human MSC transcriptome and proteome. We found that senescent MSCs displayed an aberrant response to changing substrate stiffness concomitant to their characteristic impaired differentiation potential. Furthermore, transcript and protein levels associated with the MSC intracellular proteome, the matrix and SASP exhibited different changes following the onset of senescence depending on substrate stiffness. Finally, unbiased analysis indicated a consistent, concerted downregulation of the cytoskeletal protein homeostasis complex, “chaperonin containing TCP-1/TCP-1 ring complex” (CCT/TRiC) in senescent MSCs. Chemical inhibition of CCT/TRiC activity was found to replicate the effects of senescence on mechanically-induced MSC morphology phenotypes, suggesting a role for in CCT/TRiC in the maintenance of mechanotransduction signalling pathways.

## RESULTS

### Replicative senescence in primary human MSCs affects mechanosensitive morphological changes

To characterise how senescence affected the cellular response to substrate stiffness, we used high-throughput morphometric analysis to compare proliferative, early passage (EP) primary human MSCs to donor-matched cells passaged to a point of replicative senescence (late passage, LP). Senescence was first demonstrated in the LP cells by positive β-galactosidase staining (Supplementary Figs. S1A, B) and loss of nuclear envelope protein lamin B1 (LMNB1) (Supplementary Figs. S1C-E) (Shimi *et al*, 2011). Furthermore, we confirmed the capacity of EP MSCs to undergo chemically-induced adipogenesis and osteogenesis when cultured on tissue culture plastic, showing also that this potential was attenuated by senescence (Supplementary Figs. S1F, G).

EP and LP MSCs were cultured for three days on soft (2 kPa) or stiff (25 kPa) collagen-I coated polyacrylamide (PA) hydrogels before fixation and imaging of nuclei (with DAPI staining) and filamentous actin (phalloidin staining; Fig. 1B). Substrate stiffnesses were chosen to reflect the mechanical diversity of the bone microenvironment, from marrow and adipose tissue (soft) to precalcified bone (stiff) (Ivanovska *et al*., 2017). We found the spread area of EP MSCs to be significantly greater on stiff vs. soft substrates (Fig. 1C and Supplementary Fig. S1H), accompanied by more prominent actomyosin stress fibres. These observations were consistent with previous reports (Engler *et al*., 2006; Swift *et al*., 2013). LP MSCs had significantly greater spread areas than EP cells on both soft and stiff substrates. However, a maintained responsiveness to stiffness was evident as LP cell area was significantly greater on the stiffer substrate. Concomitant to increased cell spreading on stiff substrates, the nuclei of EP cells had significantly increased nuclear areas (Fig. 1D and Supplementary Fig. S1I). The correlated behaviours of cellular and nuclear morphologies has previously been interpreted as reflective of interconnected structures in the cytoskeleton and nucleoskeleton, mediated by the linker of nucleoskeleton and cytoskeleton (LINC) complex of proteins that spans the nuclear envelope (Aureille *et al*, 2017; Gilbert *et al*, 2019). Disruptions to the LINC complex, for example by knockdown of constituent proteins, have been shown to affect mechanotransduction signalling to the nucleus (Tajik *et al*., 2016). Here we found that in LP MSCs, despite an increase in cell areas between stiff and soft substrates, the nuclei were actually significantly smaller. The correlated relationship between cell and nuclear areas (Buxboim *et al*, 2017), observed in EP cells but lost in LP cells, was evident when comparing cells from different primary donors (Fig. 1E). We also observed that alignment between cell and nuclear orientation was altered in LP cells on stiff substrates (Fig. 1F), again suggestive of lost connectivity between the cytoskeleton and nucleus. Taken together, these results indicate that senescence altered the cellular response to changes in stiffness, and in particular how these mechanical signals were manifested in nuclear morphology.

### Senescence affects the responsiveness of the transcriptome to substrate stiffness

To systemically interrogate the perturbed mechanoresponse observed in LP MSCs we performed transcriptomics on EP and LP MSCs grown on soft (2 kPa) and stiff (25 kPa) PA hydrogels for 4 days. A principal component analysis (PCA) of all samples showed a greater separation between EP and LP MSCs compared to growth on 2 and 25 kPa hydrogels (Supplementary Fig. S2A). Analysis of transcript fold-changes observed when comparing EP and LP MSCs grown on stiff substrates presented a narrower histogram of fold-changes compared to the same comparison on soft substrates (Fig. 2A), suggesting greater similarity between EP and LP cells when they are cultured on a stiffer substrate. Major metabolic remodeling during senescence leads to the secretion of a characteristic profile of proteins collectively known as the senescence-associated secretory proteome/phenotype (SASP) (Burton & Faragher, 2015). SASP transcriptome (as determined by the SenMayo gene set, Saul *et al*, 2022) changes – which were largely conserved between different substrate stiffnesses – were predominantly increased in LP MSCs (Supplementary Fig. S2B), further supporting the evidence that our LP MSCs were senescent. Enrichment analysis of transcriptome fold-changes showed that, as expected, cell cycle (GO:0007049) Gene Ontology (GO) biological processes were significantly downregulated with senescence (Fig. 2B), regardless of substrate stiffness. Senescence transcriptomee fold changes on soft and stiff were significantly correlated (Pearson’s R = 0.63, *p* < 0.0001), indicating that the majority of senescence-associated transcriptome changes were conserved regardless of substrate stiffness (Fig. 2C).

**Figure 2.**
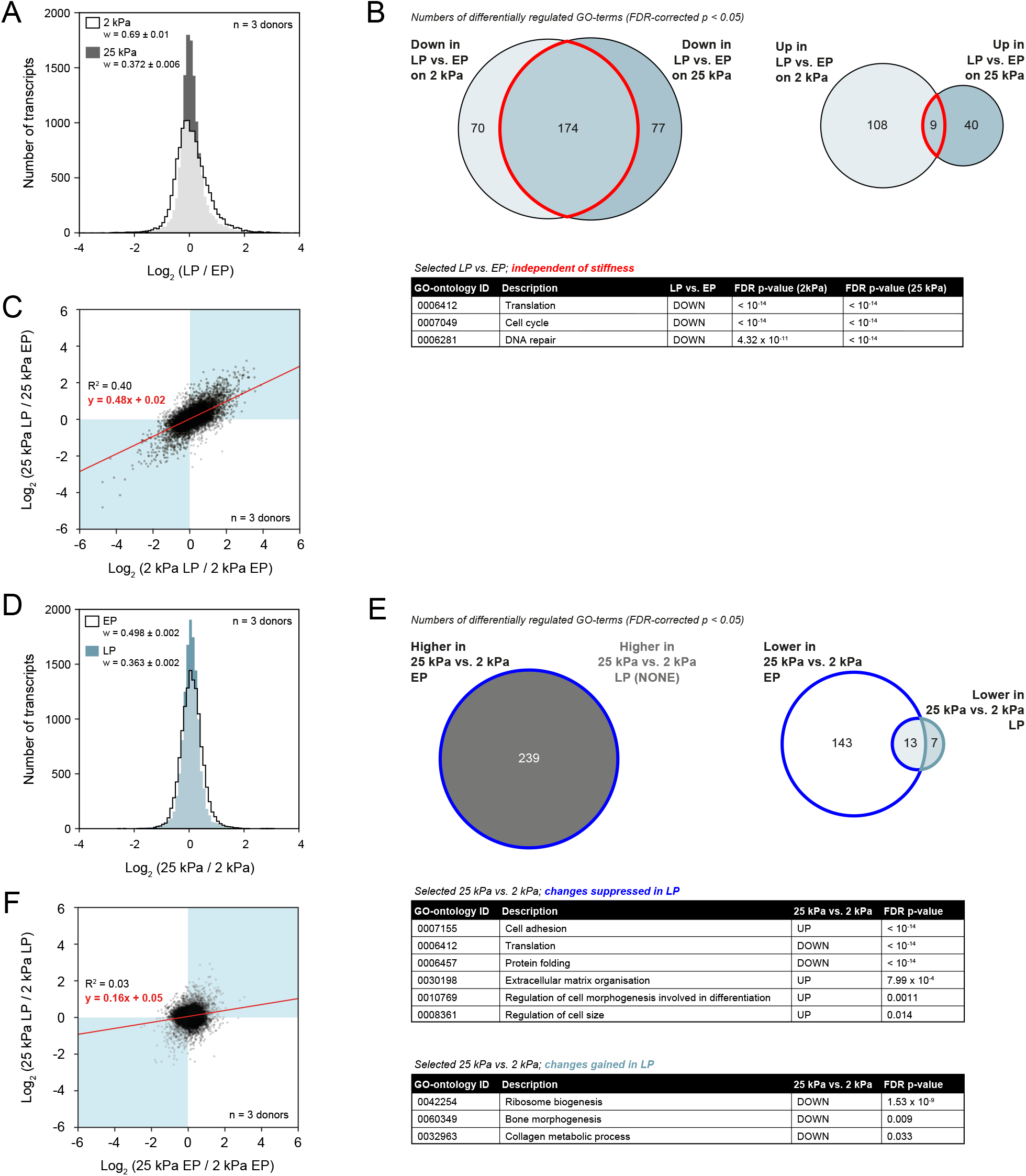
Transcriptional analysis shows an altered response to substrate stiffness in senescent cells. Proliferating early passage (EP) and senescent late passage (LP) primary human mesenchymal stem cells (MSCs) were cultured on soft (2 kPa) and stiff (25 kPa) collagen-I coated polyacrylamide hydrogels for four days prior to analysis by RNA-Seq. (**A**) Overlaid histograms of log_2_ fold-changes to transcripts in LP vs. EP cells, cultured on soft (white) or stiff (grey) substrates. The width (w) of the distribution was greater for analysis of cells cultured on soft substrate. (**B**) Venn diagrams showing numbers of unique and shared significantly differentially regulated gene ontology (GO)-terms (false discovery rate, FDR-corrected *p* < 0.05) in LP vs. EP comparisons on soft and stiff substrates. The table shows selected GO-terms significantly down-regulated in LP vs. EP cells irrespective of substrate stiffness (regions of Venn diagrams bounded in red). (**C**) Correlation plot between LP vs. EP changes in transcript on stiff substrate, compared to those on soft substrate. Senescence-induced changes to the transcriptome were correlated on soft and stiff substrates (indicated by Pearson’s R^2^ = 0.40), although the overall magnitude of the response was lower on the stiff substrate (indicated by slope of red line < 1). (**D**) Overlaid histograms of log_2_ fold-changes to transcripts in MSCs cultured on stiff vs. soft substrates; comparisons between EP cells are shown in white, LP cells in blue. (**E**) Venn diagrams showing numbers of unique and shared significantly differentially regulated GO-terms (FDR-corrected *p* < 0.05) in EP and LP MSCs cultured on stiff vs. soft substrates. Tables show selected GO-terms that were significantly differentially regulated with changing substrate stiffness in EP MSCs (top table, regions of Venn diagrams bounded by dark blue) and LP MSCs (bottom table, region of Venn diagram bounded by light blue). (**F**) Correlation plot between stiff vs. soft changes in transcript, comparing LP to EP cells. Correlation was poor (Pearson’s R^2^ = 0.03), suggesting that the mechanoresponse was not well conserved in senescent cells. All analysis based on comparisons between donor-matched cells, *n* = 3 donors.

Comparison of soft vs. stiff in EP and LP MSCs produced qualitatively similar histograms, suggesting that both EP and LP MSCs were responsive to changes in substrate stiffness (Fig. 2D). However, a subsequent analysis of enriched GO: biological process terms showed that EP and LP MSCs responded very differently to changes in substrate stiffness, with very few terms shared between EP and LP responses to stiffness (Fig. 2E). GO terms significantly enriched in stiff vs. soft in EP MSCs included downregulation of translation and protein folding, and up-regulation of cell adhesion, extracellular matrix (ECM) organization and processes involved in regulation of cell size (as expected given the increase in cell size observed in Fig. 1) and differentiation (Fig. 2E, *changes suppressed in LP*). In contrast, transcriptomes of LP MSCs grown on soft vs. stiff substrates showed significant enrichment for down-regulation of ribosome biogenesis, bone morphogenesis and collagen metabolism (Fig. 2E, *changes gained in LP*). Qualitatively, LP transcriptomes appeared responsive to changing substrate stiffness (Fig. 2D). However, correlation of EP and LP MSC transcript response to substrate stiffness showed no correlation (Pearson’s R^2^ = 0.03), suggesting a greatly perturbed mechanoresponse in LP MSCs that did not match the response of EP MSCs (Fig. 2F). Indeed, we also observed senescence-associated perturbation in the stiffness-directed response of gene targets of the mechanosensitive YAP/TAZ pathway (identified using the widely used Cordenonsi_YAP_conserved gene set, Cordenonsi *et al*, 2011). Specifically, YAP/TAZ targets were mostly increased in EP MSCs with 7 (*ITGB5, AXL, FLNA, AGFG2, TNS1, ITGB2* and *AMOTL2*) out of 51 reaching significance (Supplementary Fig. S2C, *black bars*). However, many of these fold changes were reversed in direction by senescence and only 2 (*CRIM1* and *SCHIP1*) out 51 were significantly increased (Supplementary Fig. S2C, *blue bars*). Overall, these results suggest that transcriptomic changes elicited by senescence alter known mechanosensitive pathways such that LP MSCs aberrantly respond to changing substrate stiffness like EP MSCs.

### Senescence alters the mechano-responsiveness of cell-secreted matrix components

To characterize whether senescence-associated changes to matrix composition could cause an aberrant mechanoresponse we employed label-free mass spectrometry proteomic analysis to interrogate how senescence affected the composition of cell-secreted matrix (Fig. 3A). In contrast to MSC transcriptomes which separated predominantly by LP vs EP (Supplementary Fig. S2A), a PCA of normalised peptide intensities from extracted matrix proteins showed a clearer separation of samples by substrate stiffness than by senescence (Fig. 3B). Conserved (i.e., independent of substrate stiffness) senescence-associated changes included significant decreases in type-I collagens (COL1A1, COL1A2), type-VI collagens (COL6A1, COL6A3), along with matrix proteins such as fibronectin 1 (FN1), elastic fibulin 2 (FBLN2) and tenascin-C (TNC) (Figs. 3C, D). Notably, we found that we were able to recapitulate the changes in FN1, FBLN2 and TNC by treating EP MSCs with either of the contractility-inhibiting drugs Y-27632 (ROCK inhibitor; Supplementary Fig. S3A) or blebbistatin (myosin-II inhibitor; Supplementary Fig. S3B). Furthermore, while both drugs elicited their own unique matrix proteome responses, the shared response of FN1, FBLN2 and TNC indicates that there are common effects of inhibited cell contractility that manifest in the cell-secreted matrix (Supplementary Figs. S3C). Importantly, these results suggest that some of the matrix changes observed in senescence may be due to perturbations of cell contractility.

**Figure 3.**
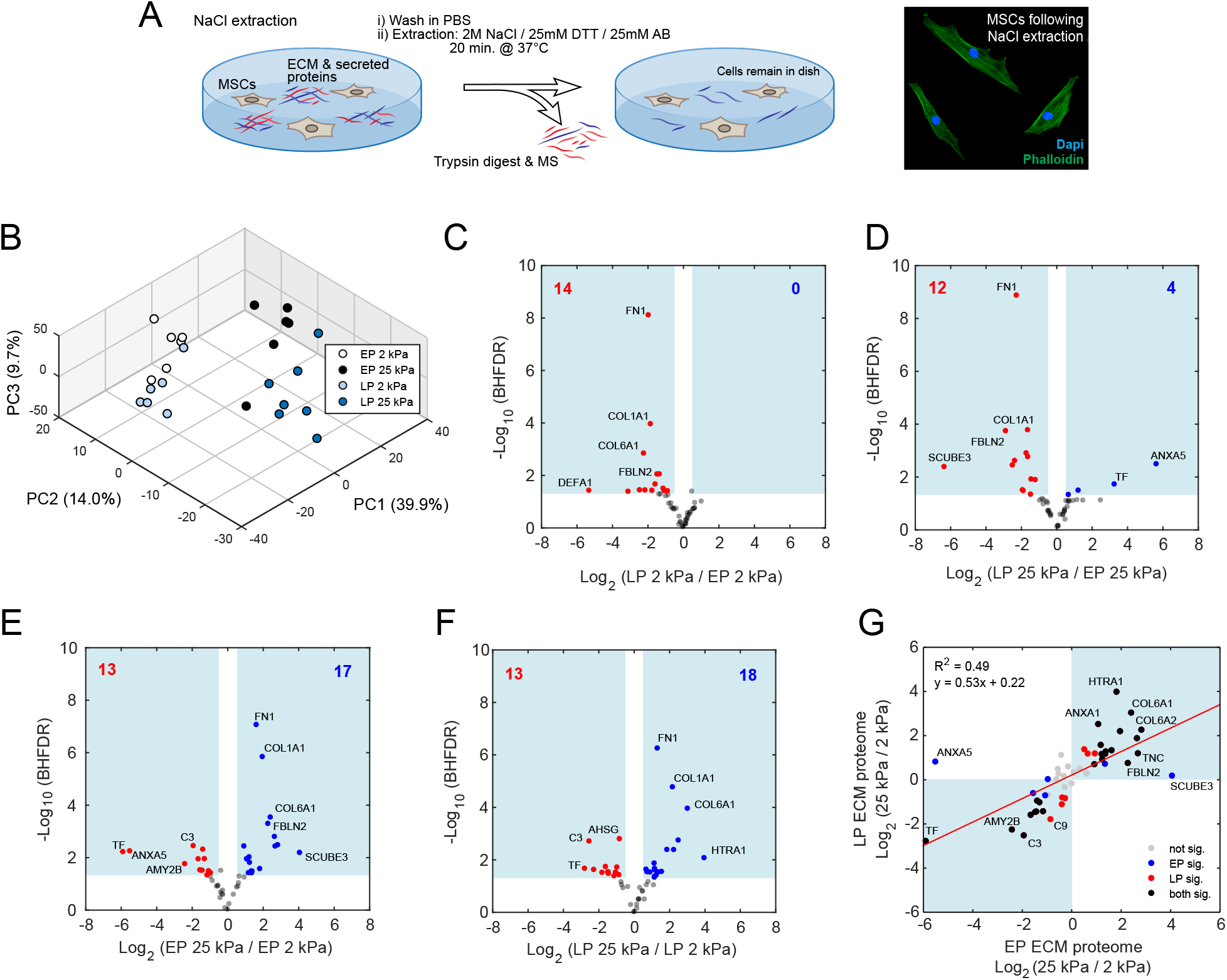
Senescence and substrate stiffness affect the cell-secreted matrix. (**A**) Schematic diagram showing how cell-secreted extracellular matrix (ECM) proteins were extracted from early passage (EP) and late passage (LP) primary human mesenchymal stem cells (MSCs) cultured on soft (2 kPa) and stiff (25 kPa) hydrogel substrates using a high ionic strength extraction buffer. The extraction buffer, which contained 2 M NaCl and the reducing agent dithiothretol (DTT) in an ammonium bicarbonate (AB) buffer, solubilised matrix proteins for analysis by mass spectrometry (MS) proteomics, while leaving cellular structures intact (image). **(B**) Principal component analysis (PCA) of donor-normalized peptide intensities following MS analysis of matrix from EP and LP MSCs, cultured on soft and stiff hydrogels. Separation in the two largest principal components (PC1 and PC2) was greater between different stiffnesses than between EP and LP. Volcano plots showing matrix proteins differentially regulated in, (**C**) LP vs. EP cells on soft substrate; (**D**) LP vs. EP cells on stiff substrate; (**E**) EP cells cultured on stiff vs. soft substrate; (**F**) LP cells cultured on stiff vs. soft substrate. In panels (C) through (F), blue and red numbers indicate how many matrix proteins were significantly up or down regulated, respectively (|log_2_ fold-change| > 0.5; Benjamini-Hochberg false discovery rate (BHFDR) adjusted *p*-value < 0.05). (**G**) Comparison of the matrix proteome response to changing substrate stiffness in EP and LP MSCs. Slope of the resulting regression line shows that mechanoresponse in LP MSCs was broadly conserved (Pearson’s R^2^ = 0.49), but displayed an overall diminished response to substrate stiffness compared to EP MSCs (slope of red line < 1). All analysis based on comparisons between donor-matched cells, *n* = 6 donors.

To interrogate how these senescence-associated matrix changes may affect the MSC mechanoresponse, we performed the corollary comparison, looking at changes in cell-secreted matrix in response to substrate stiffness in EP and LP MSCs. Analysis of the response to changing substrate stiffness showed that both EP (Fig. 3E) and LP (Fig. 3F) MSCs were capable of significantly regulating their matrix proteome as a function of substrate stiffness. This included an upregulation with increasing substrate stiffness of many of the mechanosensitive matrix proteins we had identified earlier, such as COL1, COL6, FN1, FBLN2 and TNC. However, regression analysis of the observed fold-changes showed that, on average, LP MSCs possessed a decreased magnitude of stiffness responsiveness compared to EP MSCs (Fig. 3G). Overall, these results showed that cell-secreted matrix composition was determined by substrate stiffness, but that LP MSCs – while still maintaining some capacity to respond to substrate stiffness – had a reduced ability to adapt their matrix secretion.

### Intracellular proteome of senescent MSCs reflects that of early-passage MSCs on stiff substrates

To further interrogate the cause of the perturbed matrix composition and stiffness mechanoresponse in LP MSCs, we performed label-free proteomics to analyse the changes to the intracellular proteome that occur in response to stiffness, and in senescence. Pair-wise correlation plots of peptide intensities showed a modest (Pearson’s R = 0.3 - 0.89) positive correlation between all samples following normalisation to run and donor averages (Fig. 4A), indicative of high reproducibility despite apparent donor-donor variability. In common with our transcript-level analysis (Fig. 2), volcano plots of changes in protein level with response to stiffness showed fewer significant differences in LP vs. EP MSCs (|log_2_ fold-change| > 0.5; Benjamini-Hochberg false discovery rate (BHFDR) adjusted *p*-value < 0.05; Figs. 4B, C). Correlations between transcript and protein changes were weak, potentially indicative of a lag between observed proteome and its respective transcriptome (Supplementary Figs. S4A, B). We also observed fewer significant protein fold changes between EP and LP MSCs grown on soft substrates compared to stiff substrates (Figs. 4D, E). Correlation between senescence-regulated proteome and transcriptome was weak (Supplementary Figs. S4C, D), but still stronger than that observed for stiffness-regulated transcriptome and proteome (Supplementary Figs. S4A, B). PCA of donor-normalised peptide intensities showed two distinct clusters: (1) EP MSCs on soft substrates; and, (2) EP MSCs on stiff, clustering together with LP MSCs regardless of substrate stiffness (Fig. 4F). This clustering of LP MSCs suggested that the proteomes of LP MSCs – regardless of substrate stiffness – intrinsically mirrored that of EP MSCs cultured on stiff substrates. Furthermore, this failure of LP cells to remodel their proteome in response to stiffness could provide an explanation for lost or dampened mechanosensitive regulation of the transcriptome and secreted proteins. Taken together, this multi-omic analysis has provided a comprehensive characterisation of an abrogation of the MSC mechanoresponse in senescence.

**Figure 4.**
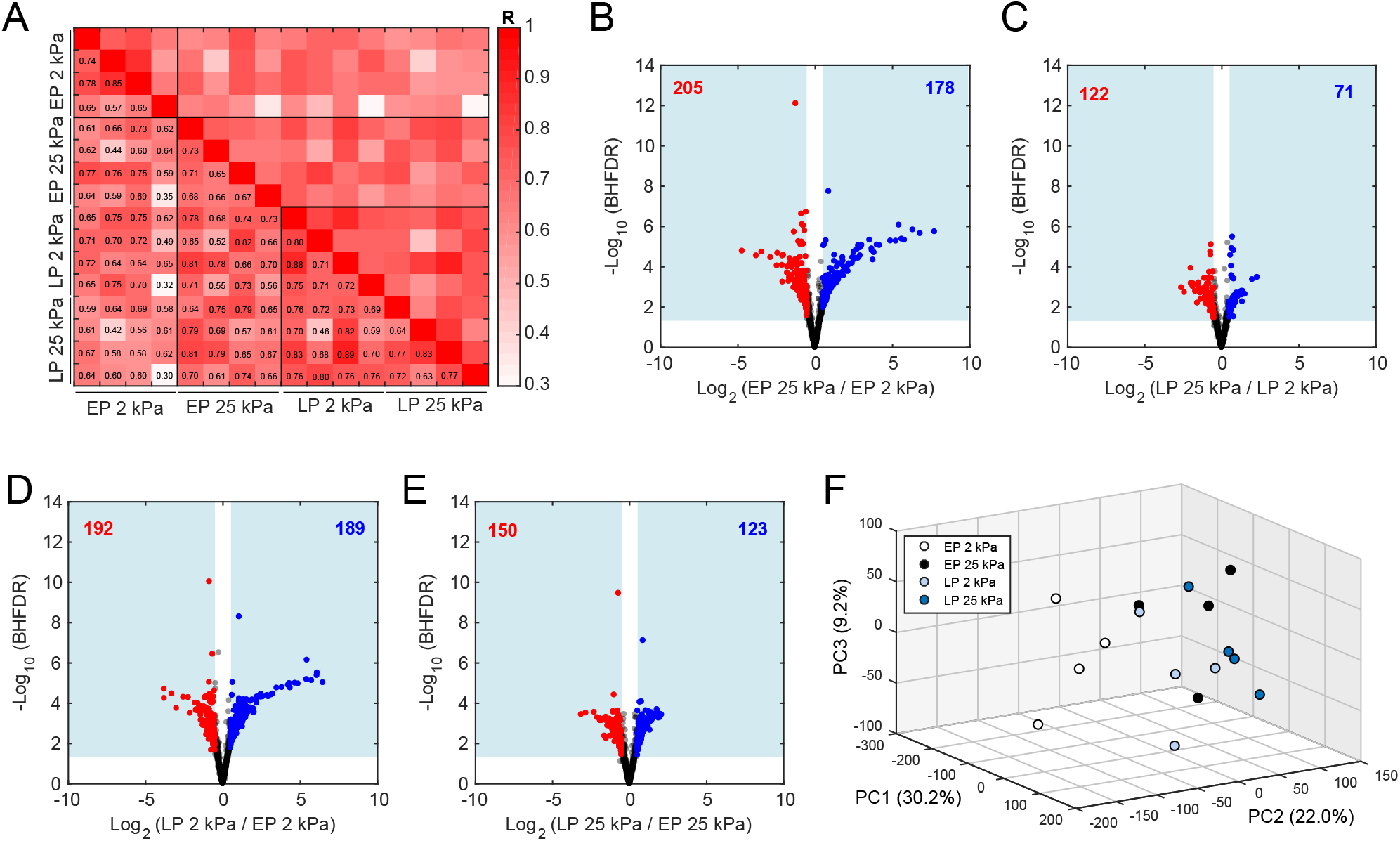
Intracellular proteomics show that cellular remodelling in response to substrate stiffness is abrogated in senescence. Mass spectrometry analysis of intracellular proteins was performed on proliferating early passage (EP) and senescent late passage (LP) primary human mesenchymal stem cells (MSCs) cultured on soft (2 kPa) and stiff (25 kPa) hydrogels. (**A**) Correlation plot of normalized intracellular peptide intensities from EP and LP MSCs cultured on soft and stiff hydrogels. Colour intensities indicate Pearson R for single pairwise comparisons. Volcano plots showing differential levels of intracellular proteins in, (**B**) EP cells on stiff vs. soft substrate; (**C**) LP cells on stiff vs. soft substrate; (**D**) LP vs. EP cells on soft substrate; (**E**) in LP vs. EP cells on stiff substrate. In panels (B) through (E), blue and red numbers indicate how many proteins were significantly up or down regulated, respectively (|log_2_ fold-change| > 0.5; Benjamini-Hochberg false discovery rate (BHFDR) adjusted *p*-value < 0.05). (**F**) Principal component analysis (PCA) plot showing clustering of samples based on donor-normalized peptide intensities. LP MSC proteomes clustered with stiff EP MSC proteomes, regardless of the actual substrate stiffness. All analysis based on comparisons between donor-matched cells, *n* = 4 donors.

### Senescent MSCs show down-regulation of the cytoskeletal chaperone complex CCT/TRiC

To interrogate why senescent MSC proteomes appeared to behave as if they were on a stiffer substrate we next sought to identify which mechanosensitive elements were altered during senescence. YAP/TAZ signalling has previously been linked to ageing and senescence in mice (Fu *et al*, 2019; Sladitschek-Martens *et al*, 2022; Xu *et al*, 2021). While we didn’t observe any significant change in *YAP1* transcript levels (Supplementary Fig. S5A), we did observe significant senescence-associated changes (mostly at the protein level) in YAP/TAZ target genes (identified using the widely used Cordenonsi_YAP_conserved gene set, Cordenonsi *et al*., 2011), on both 2 and 25 kPa hydrogels (Supplementary Fig. S5B), supporting the idea that mechanotransduction was perturbed in LP MSCs. Further analysis of cytoskeletal-associated genes and proteins (responsible for mediating cell contractility and stiffness, and identified according to the GO term “cytoskeletal organisation”, GO:0007010) showed that their abundance was mostly unaffected by senescence. However, a greater number were significantly altered during senescence on soft substrates compared to senescence on stiff substrates (Fig. 5A), consistent with the observation that senescent MSCs maintained a “stiff substrate phenotype”, irrespective of the actual substrate stiffness. A notable exception was the significant upregulation of vimentin (VIM), previously reported in senescent fibroblasts (Nishio & Inoue, 2005; Nishio *et al*, 2001), which was observed specifically on stiff hydrogels. Unbiased pathway analysis using linear modelling (PALM, Mallikarjun *et al*, 2020) highlighted a number of senescence-mediated, significantly differentially regulated pathways on both soft and stiff substrates (Fig. 5B). Senescence-mediated upregulated Reactome pathways included glycosphingolipid metabolism (R-HSA-1660662), an annotation which includes the senescence marker β-galactosidase. Senescence-mediated downregulated processes included RNA Polymerase I promoter escape (R-HSA-73772) which was represented in this dataset by a number of histone and chromatin-associated proteins. Notably, PALM analysis also highlighted significant downregulation of the cytoskeletal chaperone complex CCT/TRiC (Fig. 5B). Unlike other mentioned pathways, downregulation of CCT/TRiC components during senescence was apparent at both transcript and protein levels, regardless of substrate stiffness (Fig. 5C), suggesting concerted downregulation as a potentially integral part of MSC senescence. Indeed, CCT/TRiC components are known to be necessary for maturation of specific cell cycle proteins (Camasses *et al*, 2003; Kaisari *et al*, 2017; Liu *et al*, 2005), suggesting a potential reason for its downregulation in senescence. As CCT/TRiC is involved in folding numerous cytoskeletal components (Gao *et al*, 1992; Melki *et al*, 1997; Yaffe *et al*, 1992), we hypothesised that a loss of cytoskeletal protein homeostasis could cause a perturbation to the morphology and mechanoresponse of MSCs, as we observe in senescence.

**Figure 5.**
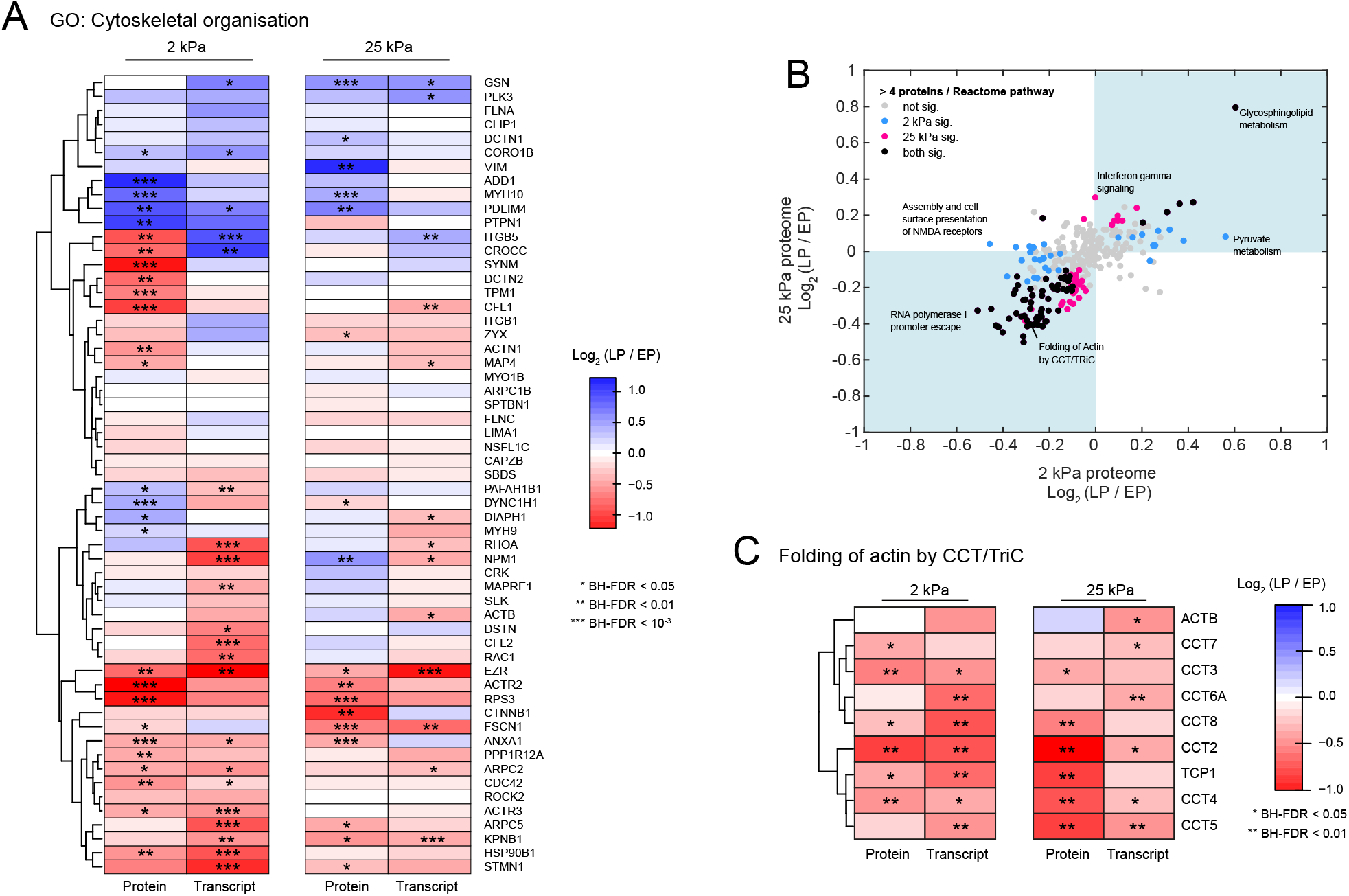
Senescence causes dysregulation of cytoskeletal proteins and downregulation of components of the cytoskeletal chaperone complex, CCT/TRiC. (**A**) Heatmap summarising intracellular protein and transcript fold-changes in senescence of primary human mesenchymal stem cells (MSCs) (i.e., comparing donor-matched late passage, LP, to early passage, EP, cells) cultured on soft (2 kPa) and stiff (25 kPa) hydrogels for genes encoding known cytoskeletal and cytoskeletal-associated proteins, as determined by the gene ontology (GO) biological process “Cytoskeletal organisation” (GO:0007010). Indicated *p*-values are corrected for Benjamini-Hochberg false discovery rate (BHFDR). (**B**) Scatter plot comparing Reactome pathway-level fold-changes elicited in MSCs by senescence on either soft (x-axis) or stiff (y-axis) hydrogels. Coloured dots indicate significant difference (BHFDR < 0.05) for that pathway in the indicated comparison. **(C)** Heatmap summarising intracellular protein and transcript fold-changes in senescence of MSCs cultures on soft and stiff hydrogels for genes belonging to the “Folding of actin by CCT/TRiC” Reactome pathway (R-HSA-390450). BHFDR-corrected *p*-values are indicated. Transcript data as in Fig. 2, *n* = 3 donors; intracellular proteomics data as in Fig. 4, *n* = 4 donors.

### Inhibition of CCT/TRiC activity in EP MSCs replicates senescent MSC morphology on soft substrates

CCT/TRiC is a highly conserved protein complex responsible for folding ∼10% of the eukaryotic proteome (Yam *et al*, 2008). CCT/TRiC is composed of two rings, each with 8 unique subunits. To interrogate the relevance of the decrease in CCT/TRiC components we observed in LP MSCs we pharmacologically inhibited CCT/TRiC using two independent, previously-characterised inhibitors: arsenic trioxide (As_2_O_3_) or the small molecule HSF1A (Neef *et al*, 2010). Arsenic trioxide has been demonstrated to inhibit CCT/TRiC (Pan *et al*, 2010) and is known to cause senescence in MSCs (Cheng *et al*, 2011) and articular chondrocytes (Chung *et al*, 2020). Studies in budding yeast have suggested that As_2_O_3_ may inhibit ATP hydrolysis by CCT/TRiC, preventing proper folding of actin substrates (Pan *et al*., 2010). Conversely, inhibition of CCT/TRiC using HSF1A does not inhibit ATP hydrolysis and has been shown to upregulate levels of the 70 kDa heat shock protein (HSP70) family, due to activation of the transcription factor heat shock factor protein 1 (HSF1) (Neef *et al*, 2014). Treatment of EP MSCs with either 1 µM As_2_O_3_ or 50 µM HSF1A for 24 hours was sufficient to significantly upregulate levels of HSP70-family member heat shock 70 kDa protein 1A (HSPA1A) specifically on soft substrates (Figs. 6A, B and Supplementary Figs. S6A, B). Culture on stiff substrates also significantly upregulated levels of HSPA1A (Figs. 6A, B). However, this stiffness-associated increase was not further enhanced by drug treatment (Figs. 6A, B), suggesting that increased stiffness may upregulate HSPA1A through inhibition of CCT/TRiC. Importantly, treatment with either CCT/TRiC inhibitor on soft substrates was sufficient to force cell and nuclear spreading (Figs. 6A, C, D and Supplementary Figs. S6C-F), a morphological phenotype similar to that observed in LP MSCs (Figs. 1B, C). These results suggest that the downregulation of CCT/TRiC that occurs during senescence in MSCs may contribute to the observed aberrant mechanoresponse.

**Figure 6.**
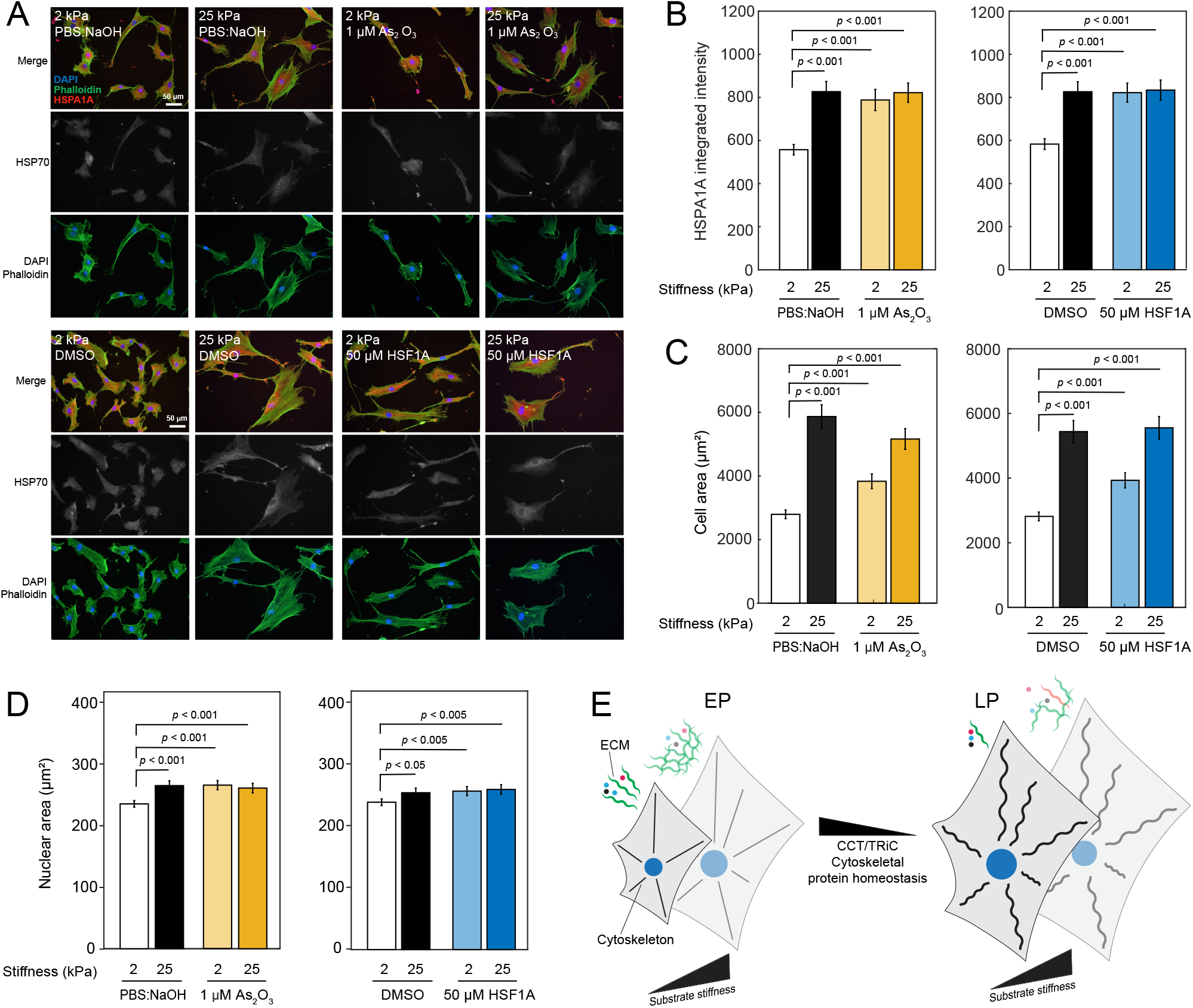
Pharmacological inhibition of the cytoskeletal chaperone complex, CCT/TRiC, causes morphological changes to early passage (EP) characteristic of senescence. (**A**) Representative images of EP human mesenchymal stem cells (MSCs) cultured on soft (2 kPa) and stiff (25 kPa) hydrogels, treated with either 1 µM As_2_O_3_ or 50 µM HSF1A, or vehicles (PBS:NaOH or 90% DMSO, respectively), for 24 hours. Cells were stained with DAPI, phalloidin and against Heat shock 70 kDa protein 1A (HSPA1A). (**B**) Quantification of HSPA1A integrated intensity. (**C**) Quantification of cell spread area. (**D**) Quantification of nuclear area. *p*-values in panels (B), (C) and (D) come from F-tests following multivariate regression modelling; *n* = 3 donors, ≥ 10 fields of view per donor, per treatment. Distribution of individual donor data is shown in Supplementary Fig. S6. (**E**) Schematic diagram summarising effects of senescence on mechanotransduction. EP MSCs possess relatively high levels of CCT/TRiC and are able to properly maintain their cytoskeleton. They show increases in both cell and nuclear area when cultured on stiff vs. soft substrates. Secreted matrix (denoted by green waved lines and different coloured circles outside the grey cell bodies) is also altered with stiffness (e.g., more fibrillar collagen is secreted, accompanied by changes in secreted signalling factors). However, CCT/TRiC levels decrease in senescence, leading to altered cytoskeletal protein homeostasis, and causing aberrant cell spreading, perturbed regulation of the transcriptome and proteome, and altered matrix secretion in response to substrate stiffness.

## DISCUSSION

Age-related matrix changes have been implicated in the failing regenerative capacity of elderly tissues. Previous studies have shown that *in vivo* ageing in mice is associated with decreased numbers of MSCs in bone-marrow aspirates; these MSCs have diminished differentiation potential and a higher incidence of senescence (Kasper *et al*, 2009). In human MSCs, increased donor age led to a greater proportion of senescent cells (β-galactosidase positive), slower proliferation, more frequent apoptosis, up-regulation of p53, p21 and apoptosis regulator BAX, and decreased osteogenic potential (Zhou *et al*, 2008). Some of these features could be reversed by inhibition of TGF-β signaling (Kawamura *et al*, 2018). Our *in vitro* senescence, primary culture system recapitulated this decrease in differentiation capacity. While senescent cells typically represent a minority of cells in a given population, even at extreme ages, the systemic effects elicited by their secreted proteome are well-documented (Young & Narita, 2009). Senescence-associated changes in matrix proteome composition have been shown to affect tissue mechanical properties (Calhoun *et al*, 2016). We have demonstrated that senescence in primary human MSCs disrupts their ability to correctly respond to changing substrate stiffness. Interestingly, inhibition of cell cycle progression has previously been shown to cause cell growth due to cytoplasmic dilution, although the link between cell cycle and fluid influx remains unexplored (Neurohr *et al*, 2019). Furthermore, previous studies have highlighted perturbed actin mobilisation during senescence in fibroblasts (Tivey *et al*, 2013; Walters *et al*, 2016). Our results suggest that impaired mechanotransduction plays a key role in mediating senescence-associated cell growth. Finally, results from pharmacological inhibition of CCT/TRiC suggest that the morphological features of senescence may be partially caused by a loss of cytoskeletal protein homeostasis (Fig. 6E).

Our results highlight the importance and utility of multi-omic characterization in teasing apart the different methods of regulation used by cells in response to different perturbations. Transcriptome PCA predominantly highlighted differences between EP and LP MSCs, possibly due to the presence of a long term (i.e., established over weeks or months) senescence programme compared to the short term (days) response to substrate stiffness, which mainly occurred at the protein level. In contrast, PCA on intracellular proteomes showed LP MSCs clustering with EP MSCs on stiffer substrates, suggesting that the intracellular proteome of LP MSCs intrinsically reflected that of a stiff cell, regardless of substrate stiffness. This clustering pattern shows that progression of the ‘core’ senescence programme was mostly independent of substrate stiffness – with senescent MSCs on both soft and stiff substrates converging on a “stiff cell” intracellular proteome. Interestingly, regulation of cell cycle progression via mechanosignalling through adhesion complex interactions has been previously observed (reviewed in Jones *et al*, 2019). Multi-omic analysis identified downregulation of cytoskeletal chaperone complex, CCT/TriC components at the protein and transcript levels as a consistent feature of senescent MSCs, regardless of substrate stiffness, and could potentially explain why senescent LP MSCs displayed an intracellular proteome that was mostly similar to EP MSCs grown on stiff substrates. This observation agrees with previous studies on MSC models of Hutchinson-Gilford progeria which showed increased cytoskeletal stiffness (Mu *et al*, 2020). These results highlight an additional – mechanical – impact that senescent cells could elicit on adjacent cells in their immediate environment, aside from the biochemical impact caused by the SASP. Indeed, “altered mechanical properties” was recently included as new hallmark of ageing (Schmauck-Medina *et al*, 2022).

Downregulation of CCT/TRiC is not specific to MSCs grown on hydrogels. Specifically, we have also observed similar CCT/TRiC downregulation in MSCs grown on tissue culture plastic (Llewellyn *et al*, 2021). Furthermore, gene expression for CCT/TRiC subunits have also been observed to be downregulated with age in human brain tissue (Brehme *et al*, 2014). Interestingly, a component of CCT/TRiC, namely the T-complex protein 1 subunit gamma (CCT3), was overexpressed in liver cancer and found to interact with the transcription factor yes-associated protein 1 (YAP1) to promote its stability by preventing ubiquitination and subsequent degradation, promoting tumorigenesis and suggesting another potential role of CCT/TRiC in regulating mechanotransduction (Liu *et al*, 2019). Furthermore, human induced pluripotent stem cells have been shown to express high levels of CCT/TRiC subunits compared to differentiated cell types and overexpression of T-complex protein 1 subunit theta (CCT8) was shown to extend lifespan in nematode worms (Noormohammadi *et al*, 2016). CCT/TRiC also regulates the stability of several key proteins involved in controlling cell cycle progression (reviewed in Wang *et al*, 2020), providing an insight as to *why* this integral chaperone complex might be downregulated in senescence (i.e., it is required to properly exit the cell cycle), with deleterious consequences for cellular mechanotransduction that don’t have noticeable effects on organismal fitness until senescent cells accumulate in old age. By altering cell secretion – and to an extent modulated by stiffness – this loss of CCT/TRiC functionality could in turn affect extracellular matrix composition, and consequently have negative effects on tissue mechanics and homeostasis in ageing.

## MATERIALS AND METHODS

### Cell culture and replicative senescence

Human mesenchymal stem cells (MSCs) were sourced from knee and hip bone marrow (male and female donors, aged 58 – 80 years) using established methodology (Strassburg *et al*, 2010). Informed written consent was obtained from donors. Experiments followed guidelines and regulations in accordance with the WMA Declaration of Helsinki and the UK Human Tissue Authority. All work was performed with approval from the NHS Health Research Authority National Research Ethics Service (approval number 10/H1013/27) and the University of Manchester. MSCs were cultured on tissue culture treated polystyrene in low-glucose DMEM with pyruvate (Thermo Fisher Scientific), 10% fetal bovine serum (Labtech) and 1% penicillin/streptomycin cocktail (Sigma-Aldrich). Cell cultures were routinely checked for mycoplasma contamination. Experiments were conducted at early passage (‘EP’; passage number 2-6) or late passage (‘LP’; passage number 6-12), with replicative senescence in LP cells defined at the passage where cell population ceased to increase. Senescence was confirmed by β-galactosidase staining, used in accordance with the manufacturer’s guidance (Cell Signaling Technology), quantified by calculating percentage of positive cells per donor based on images from bright field microscopy.

### Cell treatments

MSCs were cultured on commercially-sourced polyacrylamide gels with defined stiffness (2 and 25 kPa) and bovine collagen-I functionalization (Cell Guidance Systems). Differentiation was chemically induced in MSCs using formulations described previously (Galarza Torre *et al*, 2018), or with commercially sourced cocktails used according to the manufacturer’s instructions (StemXVivo, Bio-Techne). Adipogenesis was quantified by positive staining of lipid droplets with oil red O (Sigma-Aldrich) and osteogenesis by alkaline phosphatase activity (Sigma-Aldrich), following manufacturers’ protocols in both cases. Differentiation stains were quantified using ImageJ by calculating total stain intensity per field of view. Blebbistatin (Sigma-Aldrich) was used at 5 µM; ROCK inhibitor Y-27632 (Sigma-Aldrich) was used at 10 µM. Media and chemical treatments were replaced every four days in culture. CCT/TRiC inhibitors, As_2_O_3_ (Sigma Aldrich) and HSF1A (Axon MedChem) were used at 1 µM and 50 µM, respectively. These doses have been previously shown to inhibit mammalian CCT/TRiC (Neef *et al*., 2014; Pan *et al*., 2010). Cells were cultured for 72 hours on soft (2 kPa) or stiff (25 kPa) substrates before CCT/TRiC inhibitor treatments were added and culture was continued for a further 24 hours before fixing and imaging. A stock solution of As_2_O_3_ was made by dissolving As_2_O_3_ at 1 mM (1000x) in PBS containing 16 mM NaOH (PBS:NaOH). Stock HSF1A solutions were made at 50 mM (1000x) in 90 % DMSO/10 % MilliQ H_2_O. Cells were treated with CCT/TriC inhibitors for 24 hours, following 72 hours in culture on soft or stiff hydrogels before fixing and imaging.

### Microscopy and image analysis

Samples were prepared for microscopy using methods described previously (Gilbert *et al*., 2019). In brief, cells were fixed (4% formaldehyde, VWR International, in PBS; 10 min. at RT), washed (2 × 5 min. in PBS at RT), permeabilized (1% Triton-X, Sigma-Aldrich, in PBS; 10 min. at RT) and blocked (2% bovine serum albumin, Sigma-Aldrich, 0.25% Triton-X, in PBS; 30 min. at RT). Samples were incubated overnight at 4 °C with primary antibodies against lamin-B1 (1:500; Abcam, ab16048) or HSPA1A (1:1000; Abcam, ab181606). Samples were washed (3 × 5 min. in PBS at RT) and incubated with secondary antibodies for 1 hour at RT: AlexaFluor-488 goat anti-mouse (1:2000; ThermoFisher Scientific, A11029) or AlexaFluor-594 donkey anti-rabbit (1:2000; ThermoFisher Scientific, A21207). Samples were then washed (3 × 5 min. in PBS at RT) before staining with DAPI (1:1000; Sigma Aldrich, D9542) and AlexaFluor-488 Phalloidin (1:100; Cell Signaling Technology, #8878) at RT for 20 min. Samples were washed in PBS 3 × 5 min. before imaging. Images were taken using an Axioplan 2 microscope (Zeiss; with an EC Epiplan 10x/0.25 objective lens) and processed in ImageJ (version 2.0.0, National Institutes of Health, USA). Images subject to comparison in the same experiment had matched exposure and contrast settings. Images were corrected for background fluorescence by subtracting the mean intensity of a cell-free area from each pixel. Morphometric parameters were evaluated from images of cells cultured at low density (i.e. minimising the number of cell-cell contacts) using CellProfiler (version 2.1.1, Broad Institute, USA) (Kamentsky *et al*, 2011), with exclusion of cells that appeared mitotic. The following were evaluated: cell area; nuclear area; aspect ratio (ratio of major to minor axes of an ellipse enclosing the cell or nucleus); circularity (ratio of (4π x area) to (perimeter)^2^); and compactness (the variance of the distance between cell edge and centre, divided by cell area). Correlation between cell and nuclear orientations was also analysed.

### Secreted extracellular matrix (ECM) protein harvest

12000 MSCs were seeded per well of six-well plates (tissue culture plastic or collagen-I coated PA of defined stiffness) and cultured for 8 days. Cells were washed with PBS at RT, before ECM extraction in 500 µL 2 M NaCl, 25 mM DTT in 25 mM ammonium bicarbonate (all Fisher Scientific) for 20 min. at 37 °C. Extraction with the same buffer was performed serially in two wells. Control samples, in which ECM extraction was performed on cell-free substrates incubated with media under matched conditions for 8 days were used to create background lists of ‘false positive’, contaminant and bovine-sourced peptides in subsequent mass spectrometry (MS) analysis.

### Intracellular proteome harvest

Sample preparation of MSCs cultured on polyacrylamide gels for mass spectrometry was performed as described previously (Gilbert *et al*., 2019). Briefly, EP and LP MSCs cultured on soft (2 kPa) and stiff (25 kPa) collagen-I coated hydrogels (Cell Guidance Systems) for 4 days were washed with 2 mL PBS at RT and trypsinised at 37 °C. Detached cells were diluted 1:10 in media to saturate trypsin before pelleting by centrifugation, resuspension in cold PBS and pelleting again for storage at −80 °C. Proteins from cell pellets were solubilised by bead-beating using six 1.6 mm steel beads (Next Advance) at 4 °C in 30 µL 1.1% (w/v) sodium dodecyl sulphate (Sigma), 0.3% (w/v) sodium dodecyl sulphate (Sigma) in 25 mM ammonium bicarbonate (Fluka) supplemented with protease inhibitor cocktail (Sigma), sodium fluoride (Sigma), and sodium orthovanadate (Sigma) in de-ionised water. Bead-beating was performed in a Bullet Blender (Next Advance) at maximum speed for 2 min. The supernatant was recovered after centrifugation at 10000 rpm for 5 min. for use in the following analysis.

### Protein digest for mass spectrometry (MS)

Digest buffer (1.33 mM CaCl_2_ (Sigma), 25 mM ammonium bicarbonate) was added to 50 µL of secreted ECM extract or 25 µL of intracellular proteome extract, up to a final volume of 200 µL, in tubes containing immobilized-trypsin beads (Perfinity Biosciences). Samples were then shaken overnight at 37 °C. Protein digests were reduced (addition of 4 µL x 500 mM DTT in 25 mM ammonium bicarbonate; 10 min. shaking at 60 °C) and alkylated (addition of 12 µL x 500 mM iodoacetamide, Sigma, in AB; 30 min. shaking at RT). Next, 5 µL x 10% trifluoroacetic acid (Riedel-de Haën) in deionized water was added and organic-soluble material removed by extraction with ethyl acetate (2 × 200 µL; each addition followed by 1 min. vortexing and removal of the organic layer). Peptides were then desalted using POROS R3 beads in accordance with the manufacturer’s protocol (Thermo Fisher); samples were then lyophilized. Prior to analysis by MS, samples were diluted to 300 ng/µL (for the Orbitrap Elite) or 200 ng/µL (for the Q Exactive HF) in 5% HPLC grade acetonitrile (Fisher Scientific), 0.1% trifluoroacetic acid in deionized water. Peptide concentrations were measured using a Direct Detect spectrophotometer (Millipore).

### Mass spectrometry (MS) proteomics and data processing

Digested samples were analysed using an UltiMate® 3000 Rapid Separation liquid chromatography system (RSLC, Dionex Corporation) fitted with a 75 mm x 250 µm inner diameter 1.7 µM CSH C18 analytical column (Waters). This was coupled to either an Orbitrap Elite (Thermo Fisher Scientific) mass spectrometer or a Q Exactive HF (Thermo Fisher Scientific) mass spectrometer. For the Orbitrap Elite, peptides were separated using a gradient from 92% A (0.1% formic acid, FA, Sigma, in deionized water) and 8% B (0.1% FA in acetonitrile) to 33% B, in 104 min. at 300 nL/min. For the Q Exactive HF, peptides were separated in a gradient of 95% A and 5% B to 7% B at 1 min., 18% B at 58 min., 27% B in 72 min. and 60% B at 74 min. at 300 nL/min. Samples subject to comparison were analysed in series on the same instrument, however sample order was randomised to minimise systematic errors caused by instrument drift.

Peptides were selected for fragmentation automatically by data dependent analysis. Spectra from multiple samples were aligned using Progenesis QI (version 3.0; Nonlinear Dynamics) and searched using Mascot (server version 2.5.1; parser version 2.5.2.0; Matrix Science UK), against the UniProt human database (release-2016_04). The peptide database was modified to search for alkylated cysteine residues (monoisotopic mass change, 57.021 Da), oxidized methionine (15.995 Da), hydroxylation of asparagine, aspartic acid, proline or lysine (15.995 Da) and phosphorylation of serine, tyrosine and threonine (79.966 Da). A maximum of one missed cleavage was allowed. For samples run on the Orbitrap, peptide tolerance and MS/MS tolerance were set to 5 ppm and 0.5 Da, respectively. For samples run on the Q Exactive HF, peptide tolerance and MS/MS tolerance were set to 8 ppm and 0.015 Da, respectively. Peptide detection intensities were exported from Progenesis QI as Excel spreadsheets (Microsoft) for further processing. Peptides detected in cell-free control experiments were not used for quantification. Following database searching, peptide lists and their respective raw abundances were exported from Progenesis QI as a .csv file for further analysis using the Python implementation (version 2.6.5) of BayesENproteomics (Mallikarjun *et al*., 2020). BayesENproteomics fits regularized regression models to simultaneously calculate fold changes and impute missing values, considering donor variability and post-translationally modified peptides. BayesENproteomics also incorporates weights when calculating fold changes based on Benjamini-Hochberg false discovery rate (BHFDR)-adjusted (Benjamini & Hochberg, 1995) Mascot scores for each protein’s peptide (i.e., lower confidence in a peptide’s identity decreases that peptide’s influence on the final model fit). We used a peptide false discover rate (FDR) cut-off of 0.01. P-values for proteomics were calculated by Empirical Bayes-modified t-tests and BHFDR-adjusted.

### Quantification of transcript by RNA-Sequencing (RNA-Seq)

EP and LP MSCs were cultured on soft (2 kPa) or stiff (25 kPa) collagen-I coated polyacrylamide hydrogels for four days and harvested by trypsinisation, as in the preparation for intracellular proteome harvest described above. RNA was extracted from cell pellets using the RNeasy Mini kit (Qiagen) as per the manufacturer’s instructions and the quality and quantity assessed using a NanoDrop ND-1000 spectrometer (Thermo Fisher). Strand-specific RNA-Seq libraries were prepared using the TruSeq Stranded mRNA Sample Preparation kit (Illumina). Samples were analysed on an Illumina HiSeq4000 system using FastQC (Babraham Bioinformatics). 101 bp x 101 bp paired-end reads and between 24M and 124M total reads were generated per sample. Trimmomatic (Bolger *et al*, 2014) was used to remove low quality reads and contaminated barcodes. Libraries were aligned to the hg19 assembly of the human genome using Tophat-2.1.0 (Kim *et al*, 2013); matches with the best score were reported for each read. The mapped reads were counted by genes with HTSeq7 against gencode_v16.gtf. Log-transformed transcript abundances were normalized against median values (i.e. assuming that most transcripts were unchanged between experimental conditions), before fold changes were calculated by mixed-effects regression analysis using the Matlab *fitlme()* function, according to:

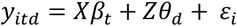

Where, *y*_*itd*_ is the normalized log-transformed abundance of transcript *i*, obtained under treatment condition *t* from donor *d*; *β*_*t*_ represents the fixed-effect logged fold changes resulting from treatment condition *t* with fixed-effects design matrix *X*; *θ*_*d*_ represents the random donor *d* effect, sampled from a multivariate Gaussian distribution with µ = 0 and σ = *ε*_*i*_(*Z*^*T*^*Z*)^−1^, *Z* is the random effects design matrix and *ε*_*i*_ is the residual variance of transcript *i*. Transcript standard error estimates and degrees of freedom were corrected using Empirical Bayes variance correction (Smyth, 2004). P-values calculated from Empirical Bayes-modified t-tests were FDR-corrected using Benjamini-Hochberg FDR (BHFDR) correction (Benjamini & Hochberg, 1995).

### Data processing and statistical treatments

Kruskal-Wallis, k-sample t-tests and sample-paired t-tests were performed in Mathematica (version 12.0; Wolfram Research Inc.) or Prism (version 8.2.1; GraphPad Software Inc.); other graphical analysis was performed in Igor Pro (version 6.37; Wavemetrics Inc.). Enrichment analysis for transcriptome data was performed using PANTHER (www.pantherdb.org), with Bonferroni correction of *p*-values enabled (Mi *et al*, 2017). Reactome (version 80) (Fabregat *et al*, 2018; Milacic *et al*, 2012) pathway analysis for intracellular proteomics data was performed using multivariate regression modelling (Mallikarjun *et al*., 2020). Significance testing for imaging data was performed using the F-tests via the *fitlm()* Matlab (version 2017a, The MathWorks) function, as in Gilbert *et al*, 2019.

## Data and code availability

Proteomics data have been deposited to the ProteomeXchange Consortium via the PRIDE partner repository with the identifiers: PXD013239 (MSCs on soft/stiff substrates treated with ROCK or myosin II inhibitors); PXD013240 (intracellular proteome extract from EP/LP MSCs grown on soft/stiff hydrogels for 4 days); and PXD013241 (secreted proteins extracted from EP/LP MSCs grown on soft/stiff substrates for 8 days). RNA-Seq data is available via EMBL-EBI ArrayExpress with identifier E-MTAB-8430. The code used to process MS data is available to download from: www.github.com/VenkMallikarjun/BayesENproteomics

## ACKNOWLEDGEMENTS

VM was partially supported by a grant from the Sir Richard Stapley Educational Trust. OD was supported by a Wellcome Trust Institutional Strategic Support Fund (097820/Z/11/B). HTJG and JS were funded by a Biotechnology and Biological Sciences Research Council (BBSRC) David Phillips Fellowship (BB/L024551/1). JL was supported by a Wellcome Trust Quantitative and Biophysical Biology studentship. Mass spectrometry was carried out at the Wellcome Centre for Cell-Matrix Research (WCCMR; 203128/Z/16/Z) Biological Mass Spectrometry Core Facility; RNA-Seq was performed by the Genomic Technologies Core Facility (GTCF) at the University of Manchester. We thank Professor Ilaria Bellantuono (University of Sheffield) and Professor Simon Hubbard (University of Manchester) for constructive criticism on the manuscript; Drs. Ronan O’Cualain, Stacey Warwood, David Knight (MS), Ping Wang, Andrew Hayes (RNA-Seq), Craig Lawless, Julian Selley (bioinformatics), Steven Marsden, Roger Meadows and Peter March (bioimaging) for Core Facility support.

## AUTHOR CONTRIBUTIONS

Investigation, VM, OD, MRJ, MK, HTJG, JL and JS; Formal Analysis, VM and JS; Writing – Original Draft, VM, JS; Writing – Review & Editing, VM, OD, MRJ, MK, HTJG, SMR and JS; Visualization, Project Administration and Funding Acquisition, JS.

