## Supplementary Information for "Loss of cytoskeletal proteostasis links dysregulation of cell size and mechanotransduction in mesenchymal stem cell senescence"

**CONTENTS**

Supplementary Figures S1 – S6 … … … … … … … Page 3

**Supplementary Figure S1.**

A

| Donor ID | Donor Age | Donor Sex | Location | EP Passage # | EP % β-gal. +ve | LP Passage # | LP % β-gal. +ve |
|---|---|---|---|---|---|---|---|
| d176 | 58 | F | knee | 3 | 24.3 | 8 | 87.5 |
| d191 | 61 | F | hip | 3 | 29.6 | 10 | 72.5 |
| d191 | 61 | F | hip | 5 | 2.6 | 12 | 57.6 |
| d312 | 62 | M | knee | 3 | 38.5 | 11 | 55.7 |
| d312 | 62 | M | knee | 5 | 0.0 | 12 | 5.3 |
| dW176 | 64 | F | hip | 3 | 31.7 | 6 | 94.7 |
| d309 | 69 | F | knee | 3 | 40.9 | 10 | 73.5 |
| d309 | 69 | F | knee | 5 | 0.0 | 12 | 12.9 |
| d112 | 74 | F | hip | 3 | 12.7 | 8 | 86.7 |
| d112 | 74 | F | hip | 6 | 3.8 | 11 | 80.4 |
| d210 | 80 | F | knee | 5 | 14.0 | 8 | 30.0 |

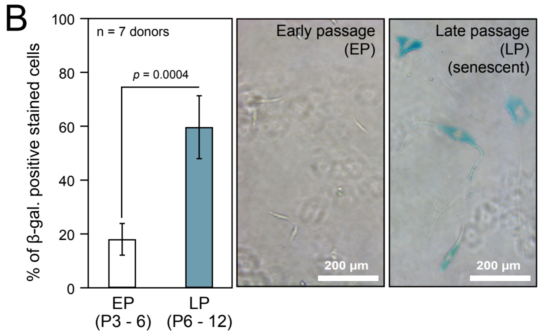


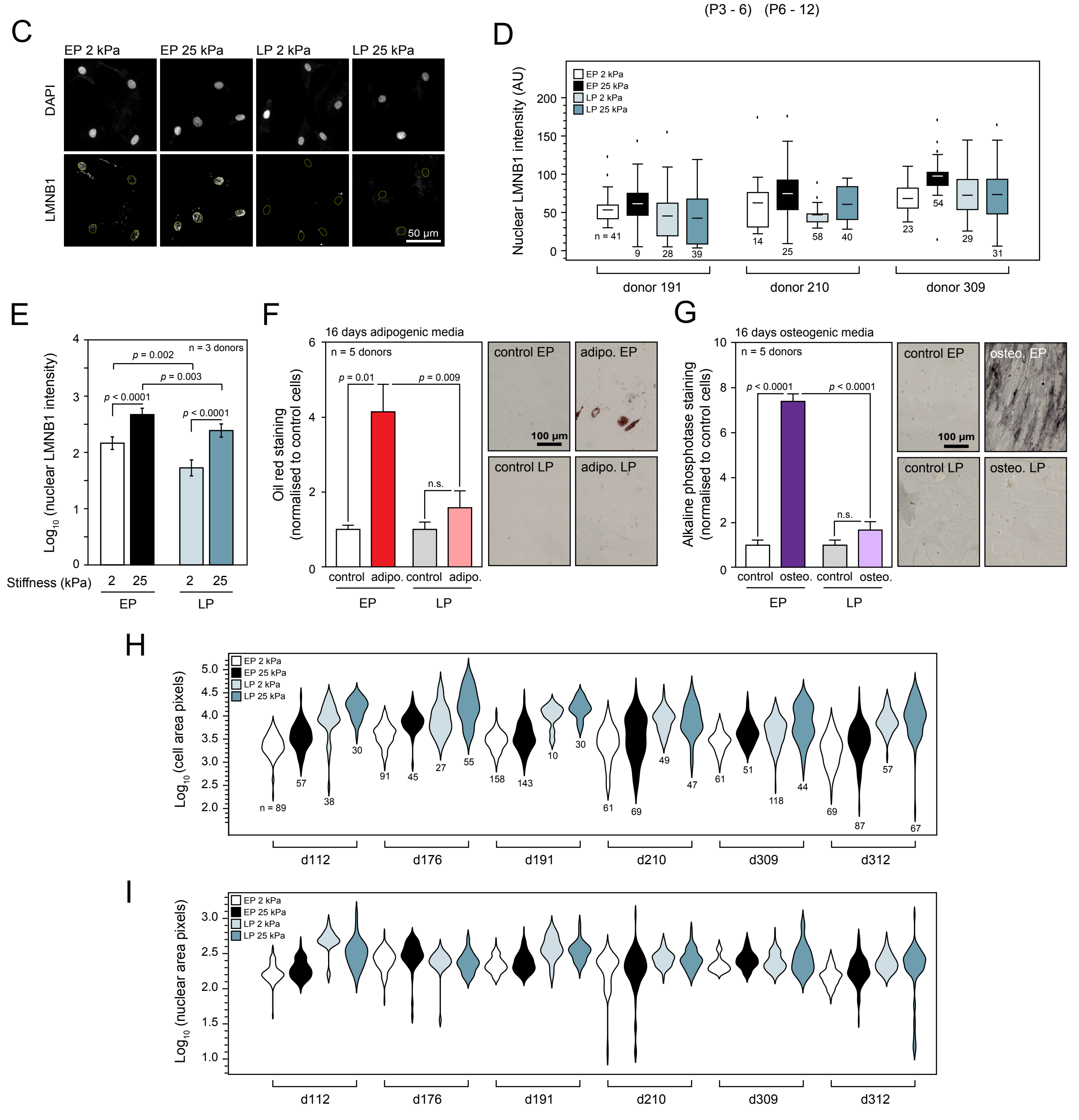

**Supplementary Figure S1. β-Galactosidase staining, lamin-B1 (LMNB1) staining and stiffness-directed morphologies of early passage (EP) and late passage (LP) primary human mesenchymal stem cells (MSCs).** (**A**) Table of donor information and the passage numbers at which MSCs were used in EP and LP states. (**B**) β-galactosidase staining was used to confirm senescence in MSCs passaged to a point where they ceased to replicate. Bars show means ± SEM; *p*-value from donor-paired t-test, *n* = 7 donors. (**C**) Representative images showing nuclear staining of DAPI and LMNB1. (**D**) LMNB1 nuclear staining quantified by donor. (**E**) Summary of quantification of nuclear LMNB1 intensities. P-values from F-tests, *n* = 3 donors. (**F**) Quantification of chemically-induced adipogenesis by Oil Red staining of EP and LP MSCs cultured for 16 days on tissue culture plastic. Adipogenic potential was significantly suppressed in senescence ($p$ = 0.009). **G**) Quantification of chemically-induced osteogenesis by alkaline phosphatase staining of EP and LP MSCs cultured for 16 days on tissue culture plastic. Osteogenic potential was significantly suppressed in senescence ($p$ < 0.0001). In panels (F) and (G), bars show means ± SEM; *p*-values from Kruskal-Wallis test with Dunn's multiple comparisons, *n* = 5 donors. Plots of the distributions of (**H**) cell areas and (**I**) nuclear areas in EP and LP MSCs cultured on soft (2 kPa) or stiff (25 kPa) collagen-I coated polyacrylamide hydrogels. Data from six donors are shown; annotations indicate the number of cells and nuclei analysed under each condition.

**Supplementary Figure S2.**

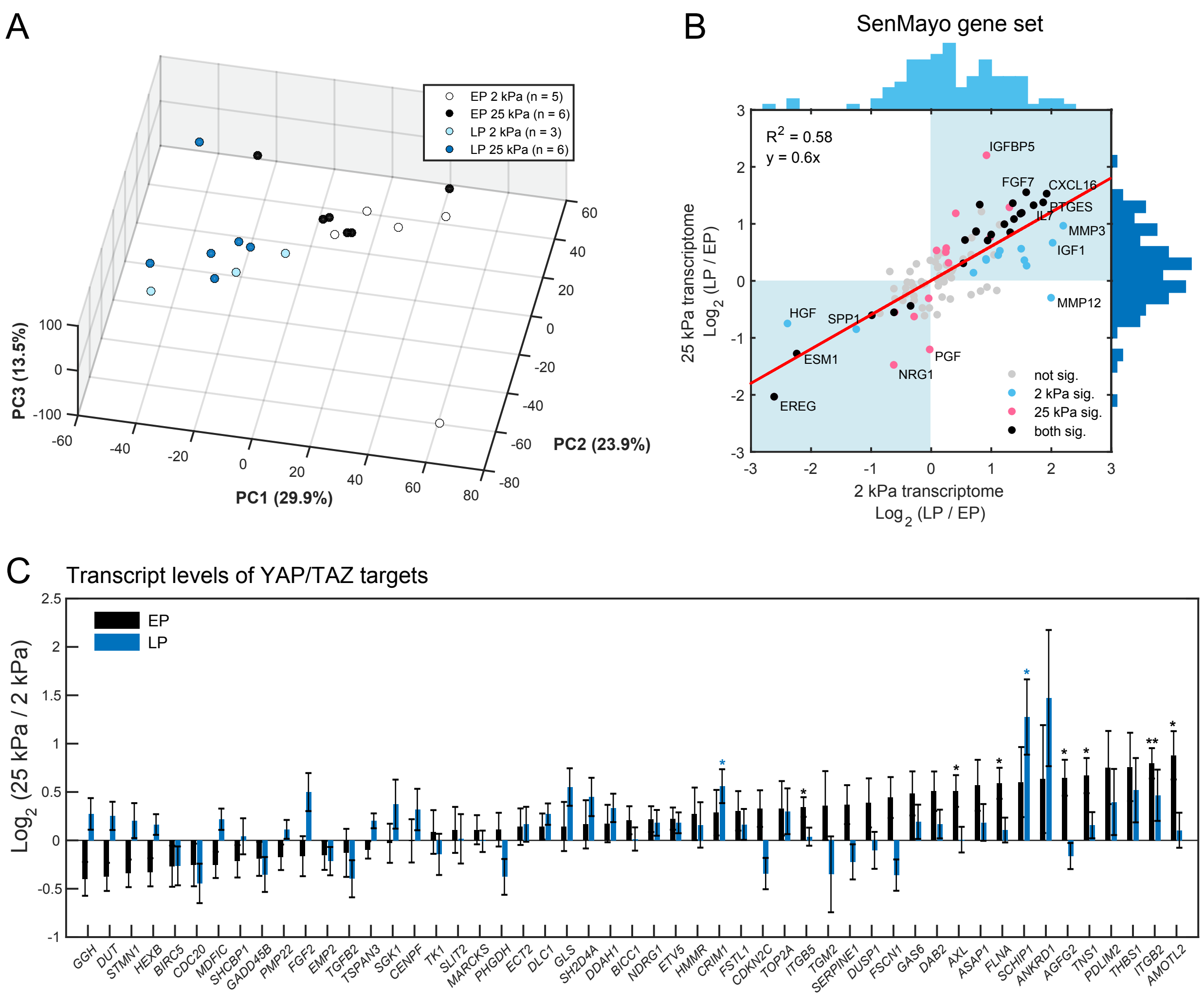

**Supplementary Figure S2. Transcriptomic analysis of early passage (EP) and late passage (LP) primary human mesenchymal stem cells (MSCs) cultured on soft (2 kPa) and stiff (25 kPa) hydrogels.** (**A**) Principal component analysis (PCA) showing separation of donor-normalised transcriptomes predominantly by EP vs. LP. (**B**) Scatter plot showing $\log_2$ fold-changes in senescence associated secretory phenotype (SASP)-associated transcripts (as determined from the SenMayo gene set [Ω]) observed when comparing LP vs. EP MSCs cultured on either soft (x-axis) or stiff (y-axis) hydrogels. Statistical significance (Benjamini-Hochberg false discovery rate (BHFDR)-corrected p-value < 0.05) is denoted by colour. Histograms show relative frequencies of respective $\log_2$ fold-changes in each comparison, showing that most SASP-associated transcripts are increased in LP MSCs relative to EP MSCs. (**C**) Bar charts showing stiffness-directed (i.e., stiff vs. soft) $\log_2$ fold-changes in expression of YAP/TAZ target genes in either EP (black bars) or LP (blue) MSCs. Bars represent linear model effects sizes ± standard errors. *, BHFDR < 0.05; **, BHFDR < 0.01.

[Ω] Saul D, Kosinsky RL, Atkinson EJ, Doolittle ML, Zhang X, LeBrasseur NK, Pignolo RJ, Robbins PD, Niedernhofer LJ, Ikeno Y et al (**2021**) A new gene set identifies senescent cells and predicts senescence-associated pathways across tissues. *bioRxiv*: 2021.2012.2010.472095

**Supplementary Figure S3.**

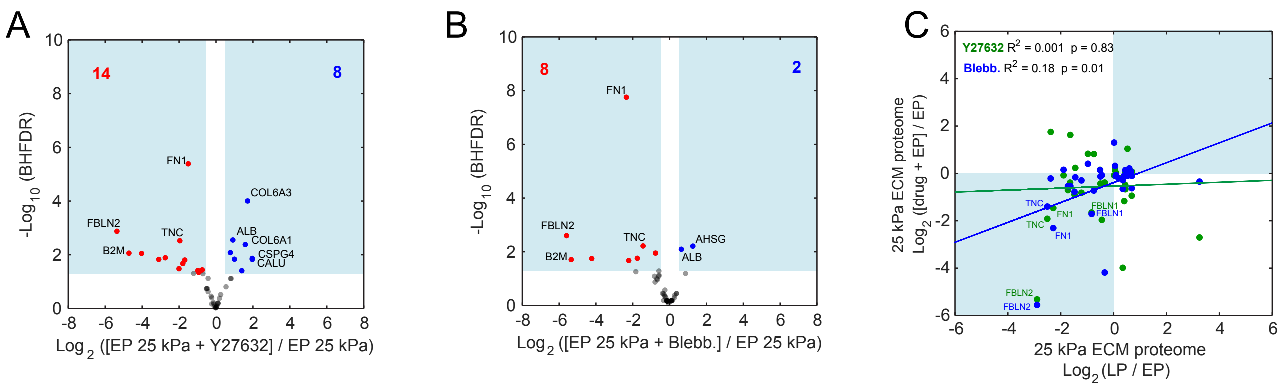


**Supplementary Figure S3. Matrix secretion from mesenchymal stem cells (MSCs) is mechanically regulated.** (**A**) and (**B**) are volcano plots showing matrix protein fold-changes (x-axis) and statistical significance (y-axis) induced by treating early passage (EP) MSCs grown on stiff (25 kPa) hydrogels with contractility inhibiting drugs, Y-27632 (**A**) or blebbistatin (**B**). Blue and red numbers indicate how many matrix proteins were significantly up or down regulated, respectively ($|\log_2$ fold-change$| > 0.5$; Benjamini-Hochberg false discovery rate (BHFDR) adjusted $p$-value $< 0.05$; $n = 3$ donors). (**C**) Scatter plot showing that perturbing cell contractility by two distinct mechanisms leads to similar changes in proteins FBLN1, FBLN2, TNC and FN1.

**Supplementary Figure S4.**

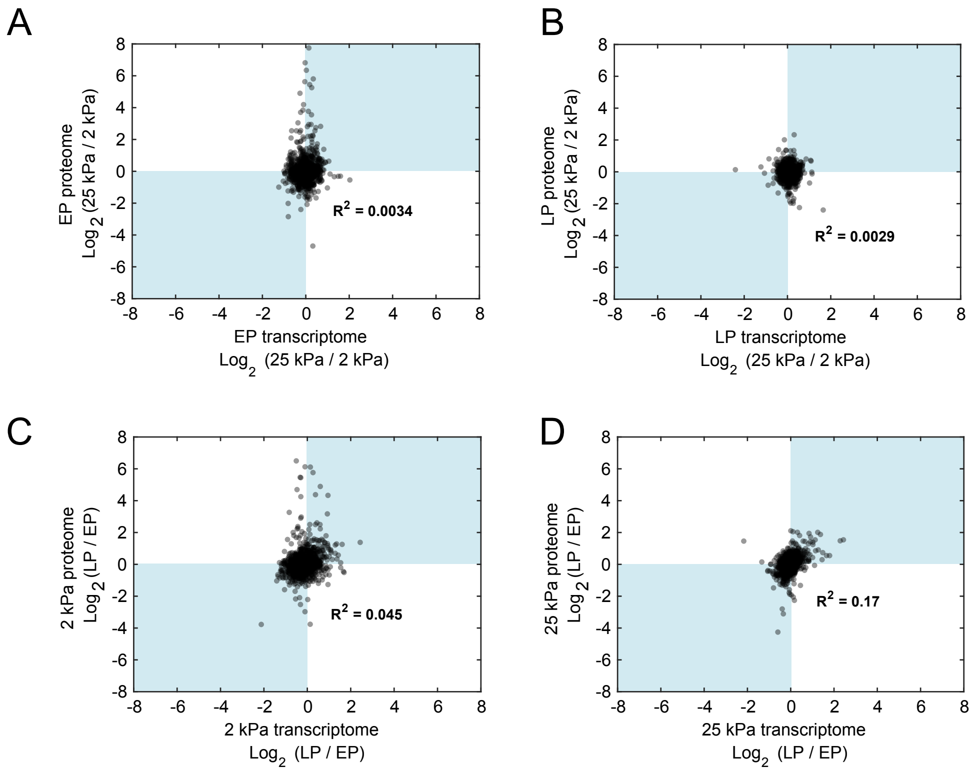


**Supplementary Figure S4. Effects of substrate stiffness and replicative senescence on intracellular proteins and transcripts.** Plotting intracellular protein and transcript fold-changes showed no correlation in both (**A**) early passage (EP) and (**B**) late passage (LP) human mesenchymal stem cells (MSCs) subjected to different substrate stiffnesses (stiff, 25 kPa vs. soft, 2 kPa). Corresponding plots of LP vs. EP protein and transcript fold-fold changes in MSCs cultured on (**C**) soft and (**D**) stiff hydrogels also showed poor correlation. Transcript data as in Fig. 2, *n* = 3 donors; intracellular proteomics data as in Fig. 4, *n* = 4 donors.

**Supplementary Figure S5.**

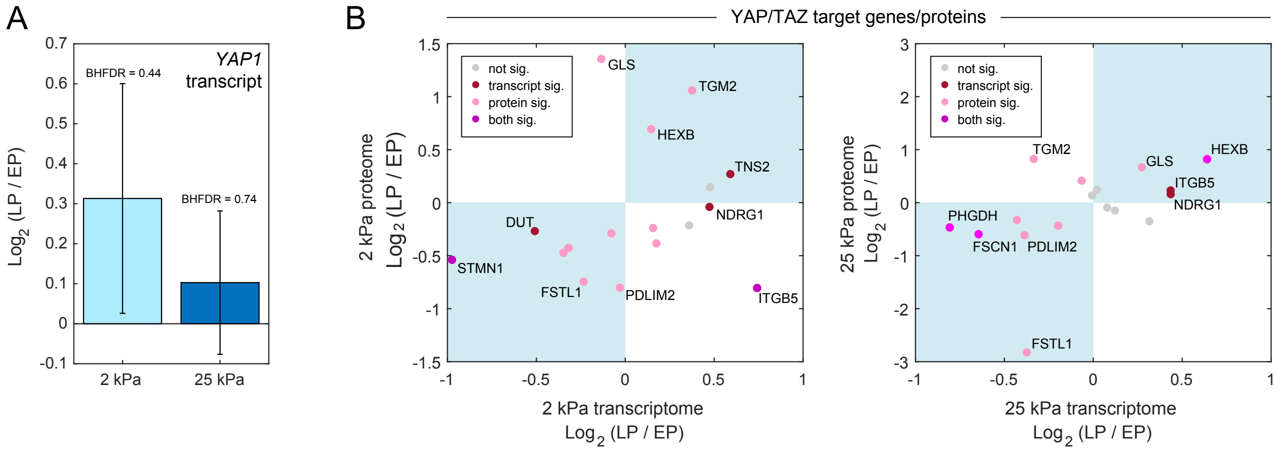


**Supplementary Figure S5. Effect of senescence on *YAP1* transcript and YAP/TAZ target genes in primary human mesenchymal stem cells (MSCs).** (**A**) Fold-changes of transcript levels of yes-associated protein 1 (*YAP1* in response to senescence (i.e., comparing late passage, LP, vs. early passage, EP MSCs) on either soft (2 kPa) or stiff (25 kPa) hydrogels, as indicated. Empirical Bayes-modified t-tests with Benjamini-Hochberg correction for multiple comparisons (indicated as BHFDR) showed no significant differences between LP and EP MSCs on either stiffness (*BHFDR* > 0.05). (**B**) Scatter plots showing logged senescence-associated (i.e., LP vs. EP) transcript (x-axis) vs. intracellular protein (y-axis) fold-changes specifically in YAP/TAZ target genes and their protein products, either on soft or stiff hydrogels. Significance at either transcript or protein-level indicated by colour and calculated by empirical Bayes-modified t-tests with Benjamini-Hochberg p-value correction for multiple comparisons, with BHFDR < 0.05 deemed significant.

**Supplementary Figure S6.**

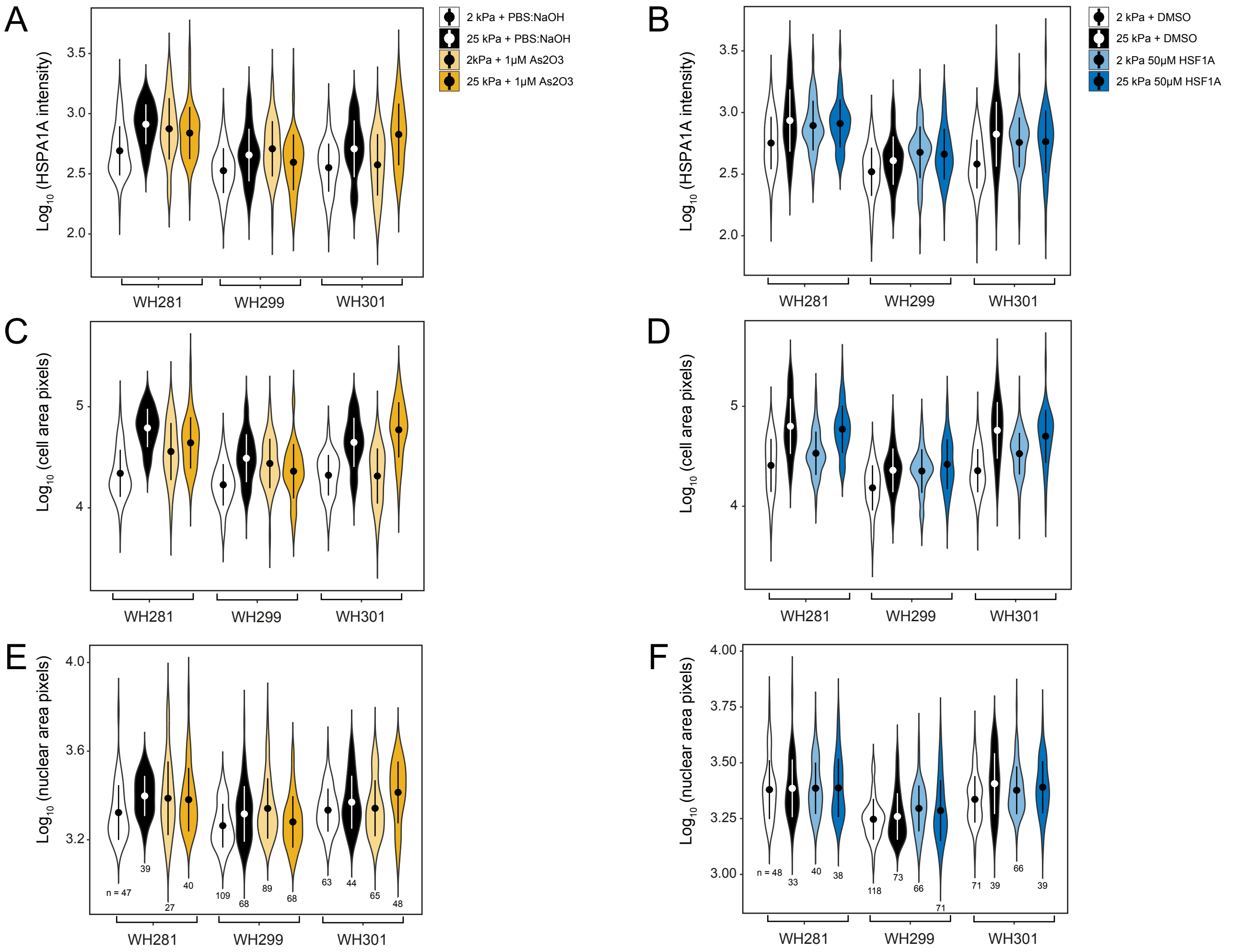


**Supplementary Figure S6. Quantification of the effects of pharmacological inhibition of CCT/TRiC activity on the morphology of primary human mesenchymal stem cells.** Violin plots showing donor-level breakdown of quantifications in Fig. 6. Summaries indicate mean ± standard deviation. (**A**) and (**B**), analysis of HSPA1A intensity; (**C**) and (**D**), analysis of cell area; (**E**) and (**F**), analysis of nuclear area (numbers indicate how many cells were analysed).
